# Surprising variety in the USP deubiquitinase catalytic mechanism

**DOI:** 10.1101/2023.07.24.550302

**Authors:** Niels Keijzer, Anu Priyanka, Yvette Stijf-Bultsma, Alexander Fish, Malte Gersch, Titia K. Sixma

## Abstract

The USP family of deubiquitinases (DUBs) controls many ubiquitin-dependent signaling events. This generates therapeutic potential, with active-site inhibitors in preclinical and clinical studies.

Understanding of the USP active site was so far primarily guided by USP7 data, where the catalytic triad consists of cysteine, histidine and a third residue (first critical residue), which polarizes the histidine through a hydrogen bond. A conserved aspartate (second critical residue) is directly adjacent to this first critical residue.

Here we study the roles of these critical residues in a subset of USPs and reveal a remarkable variety in function. While USP7 relies on the first critical residue for catalysis, this residue is dispensable in USP1, USP15, USP40 and USP48. Instead, their second critical residue is vital for catalysis.

Interestingly, without their respective vital residue USP7, USP15 and USP40 can still perform nucleophilic attack. The diverging catalytic mechanisms of USP1 and USP7 are independent of substrate and retained in cells for USP1. The unexpected variety of catalytic mechanisms in this well-conserved protein family may generate opportunities for selective targeting of individual USPs.

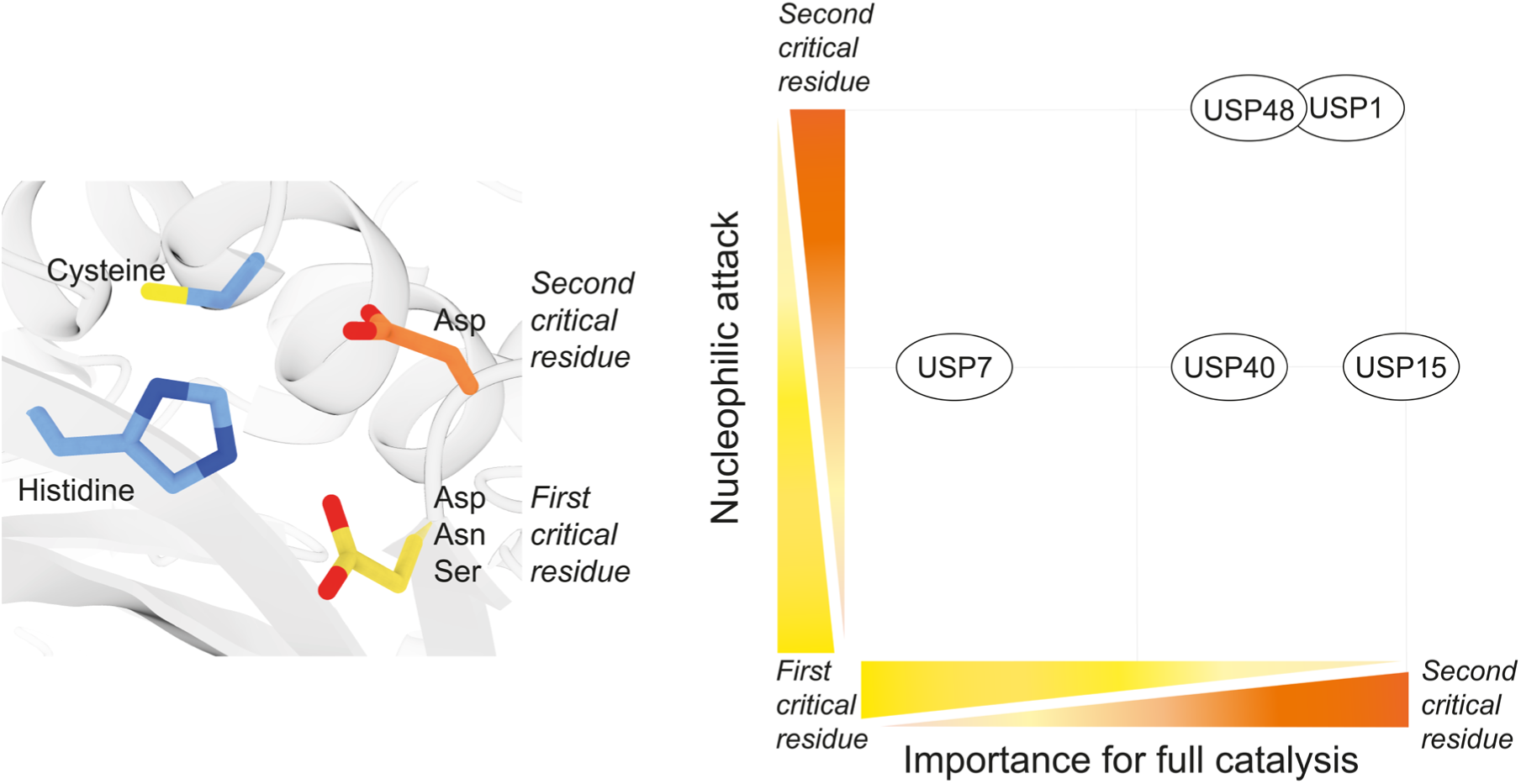

**Synopsis:**

- The roles of the highly conserved critical residues in USP active sites are poorly understood. Here we show that these two residues have varying importance for catalysis between different USPs.
- Except for USP7, the majority of USPs does not rely on the canonical third catalytic residue (first critical residue). Instead, the USPs tested rely primarily on the highly conserved second critical residue.
- In some USPs, either critical residues can accommodate nucleophilic attack (USP7, USP40, USP15). USP1 and USP48 are unable to perform the nucleophilic attack without the second critical residue.

## Introduction

Deubiquitinating enzymes (DUBs) are isopeptidases that remove ubiquitin from target substrates. This regulation of ubiquitination is essential, as it is involved in many different cellular pathways. Ubiquitination can cause changes in localization, activation, signaling of the protein, or facilitate its degradation by the proteasome or lysosome (Hershko & Ciechanover, 1998). Ubiquitin itself can be modified on its seven lysines or its N-terminal methionine, thereby generating a variety of ubiquitin chain types. These different chain types add to the vast variety in signaling potential of ubiquitin (Komander & Rape, 2012). By removing ubiquitin or ubiquitin chains from target substrates, DUBs play a vital role in regulating pathways throughout the cell and inhibition of DUBs is therefore a viable therapeutic strategy. In fact, small molecule inhibitors for USP1 and USP30 are currently being explored in clinical studies against cancer and renal disease, respectively (Cadzow et al., 2020; Tsefou et al., 2021).

Ubiquitin specific proteases (USPs) form the largest known family of deubiquitinating enzymes (Nijman et al., 2005). They were originally discovered in yeast (termed UBPs) and were identified as cysteine proteases of the papain superfamily, due to their extremely conserved cysteine and histidine residues (Baker et al., 1992; Barrett & Rawlings, 1996; Papa & Hochstrasser, 1993). Together with a third catalytic residue these form a catalytic triad similar to other proteases (Cstorer & Ménard, 1994; Stroud, 1974). In order to cleave the ubiquitin-substrate bond, the catalytic cysteine acts as the nucleophilic agent and provides the reactive thiol group. The properties of this cysteine, enabling nucleophilic attack, are endowed by a histidine (Polgár, 2013). A third catalytic residue forms a hydrogen bond with this catalytic histidine, thereby polarizing it and allowing it to stabilize the catalytic cysteine (Vernet et al., 1995). Subsequently a tetrahedral thioester intermediate is generated which is then hydrolysed to regenerate the free enzyme. Transition states for the formation and hydrolysis of the thioester are stabilized by the oxyanion hole, an essential network of hydrogen bonds provided by neighboring residues (Ménard & Storer, 1992).

All USPs contain a conserved catalytic domain (∼350 amino acids), generally decorated with additional domains that add to their extensive structural and functional variety (Nijman et al., 2005; Komander et al., 2009; Ye et al., 2009). The conformation of the USP catalytic triad was first revealed when the structure of USP7 catalytic domain was published (Hu et al., 2002). It was shown that USP7’s catalytic triad consists of cysteine, histidine, and mutagenesis identified an aspartate as the third catalytic residue. Directly adjacent to this aspartate lies another aspartate, and the structure showed that this residue plays an essential role in stabilization of the tetrahedral intermediates via a nearby water molecule (Hu et al., 2002). Following this discovery, the catalytic triad of USPs has primarily been identified using sequence and structural alignments, which show high sequence and structural conservation of all residues involved.

The residue that acts as USP7’s third catalytic residue is either an aspartate, an asparagine or a serine in other human USPs. In contrast, the succeeding residue adjacent, termed second critical residue in this work, is much better conserved, as it is almost always an aspartate in human USPs. In the context of the structural analysis of USP2’s catalytic domain it was realized that either of these two residues (N574, D575) is sufficient for catalytic activity, as mutations in either residue individually had no significant effect on catalysis (Zhang et al., 2011). However, despite this conservation, the importance and precise role of this second critical residues has not been assessed among the wider USP family. In contrast, based on structural analysis alone, a handful of papers assign the second critical residue as the catalytic residue, but do not verify it by mutagenesis (Leznicki et al., 2018; Pereira et al., 2015; Yin et al., 2015). Moreover, it is clear that the original findings in USP7 can not easily be extrapolated to other USPs (Hu et al., 2002; Davis & Simeonov, 2015) and it is not clear whether there exists a unified mechanism for this DUB family.

Several USPs harbor a misaligned catalytic triad, which was first shown in a USP7 structure (Hu et al., 2002). Upon ubiquitin binding, the switching loop (SL) of USP7 changes conformation which causes a rearrangement of the catalytic triad into its catalytically competent conformation (Faesen et al., 2011, Kim et al., 2016). USP40 has a USP7-like activation mechanism, implying a similar inactive state (Kim et al., In preparation). USP15 also harbors a misaligned catalytic triad but unlike USP7, USP15 does not require ubiquitin binding in order to realign, and instead undergoes conformational changes prior to ubiquitin binding (Priyanka et al., 2022). Although the SL is structurally well conserved among USPs, such a conformational change has only been observed in USP7, USP15, and USP34 (Xu et al., 2022). The apo-structure of USP4, which shares domain structure with USP15, did not show a misaligned catalytic triad, and neither did USP8, another USP that resembles USP15 and USP7 (Priyanka et al., 2022).

Here, we revisit the catalytic mechanism assignment of USP DUBs. We investigate a set of five diverse USPs (USP1, USP7, USP15, USP40 and USP48) and perform Michaelis Menten analysis to assess the importance of the third catalytic residue and the adjacent aspartate for catalysis. We reveal that instead of the canonical third catalytic residue, USP1, USP15, USP40 and USP48 rely on the adjacent highly conserved aspartate for catalysis. USP15 does not require the canonical third catalytic residue for catalysis, as a loss of function mutant behaves like the wildtype enzyme and USP1. USP1, USP40 and USP48 only suffer from a small decrease in activity when the canonical residue is lost. Only USP7 is rendered catalytically dead when the first critical residue is mutated. We verify the importance of the relative role of these residues by analyzing the effect of these mutations in cells for USP1. Our results demonstrate that using structural and sequence alignment alone do not predict which third catalytic residue is used and that a surprising degree of plasticity exists between the catalytic components of USPs. This finding has important implications for the dissection of catalytic mechanisms of other USPs and suggests that opportunities for selective targeting could exist.

## Methods

### Plasmids, cloning and purification

USP1/UAF1 was co-purified by co-expressing USP1 (pFastbac-HTb, N-terminal his-tag, G670A + G671A to prevent autocleavage (Huang et al., 2006)) and UAF1 (pFastbac1, N-terminal strep-tag) in Sf9 cells according to (Dharadhar et al., 2021). Codon-optimized USP7FL (pGEX-6p-1) and USP7CD (pGEX-6p-1, N-terminal GST-tag, res. 208–560) constructs (codon-optimized) (Faesen et al., 2012) were expressed in E. coli and purified following the protocol described by (Kim et al., 2016). USP15D1D2 (pET21a, C-terminal His-tag, res. 255-919, Δ440–756, codon-optimized) was expressed in E. coli and purified according to (Priyanka et al., 2022). USP40: Full length codon optimized USP40 was expressed in Sf9 cells and purified according to (Kim et al., in preparation). USP1^D751A^, USP1^D752A^, USP7^D481A^, USP7^D482A^, USP15^D879A^, USP15^D880A^, USP40^N452A^, USP40^D453A^, USP48^N370A^ and USP48^D371A^ in their corresponding vectors were generated with QuickChange site directed mutagenesis, verified by sequencing, expressed and purified analogous to wild type protein.

### Protein stability

In order to assess protein stability of all enzymes used in this study a thermal stability assay was performed using nanoDSF (Prometheus NT.48, NanoTemper Technologies GmbH). In this assay the enzymes were diluted to final concentrations of 0.5 mg/ml (USP1/UAF1, USP7) or 0.25 mg/ml (USP15, USP40 USP48). USP1/UAF1, USP7 and USP48 were tested in 20 mM Hepes, 150 mM NaCl, 5 mM DTT, 0.05% Tween-20 following previous publications (Dharadhar et al., 2021; Uckelmann et al., 2018; Faesen et al., 2011). USP15 and USP40 were tested in 20 mM Hepes pH 7.5, 100 mM NaCl, 5 mM DTT, 0.05% Tween-20 following previous publications (Kim et al., in preparation). Unfolding and aggregation of enzymes were assessed by measuring the tryptophan intrinsic fluorescence intensity over a temperature gradient from 20°C to 90°C. Using manufacturers build-in software, the melting temperatures were determined using the ratio of fluorescence intensity of 330 nm and 350 nm and onset of aggregation was determined using light scattering at 266 nm.

### Multiple sequence alignment

The sequence of USP7’s catalytic domain (residues 214-521) was used as a reference sequence to blast against the Uniprot database (Bateman et al., 2023). Catalytic domains as defined by Uniprot of the resulting human USPs were used for multiple sequence alignment. In order to properly align USP1, its inserts were removed from the catalytic domain following (Dharadhar et al., 2021). In order to properly align USP40, a shorter sequence was used (residues 250-480). Sequences were aligned using the tCoffee webserver (di Tommaso et al., 2011), and alignment was visualized using Jalview software (Waterhouse et al., 2009). A sequence logo was generated based on this multiple sequence alignment, of an 11 amino acid sequence surrounding the critical residues in a web-based sequence logo generator (Crooks et al., 2004).

### Structural superposition

Structural alignment was performed using USP catalytic domains for which a PDB structure is available (*Table 1*). Structures of USPs bound to ubiquitin were use whenever possible, to ensure a catalytically competent conformation. Structures were aligned using the ChimeraX (Goddard et al., 2018) build-in matchmaker option (Pettersen et al., 2021).

**Table 1:** List of USPs used in superposition, corresponding PDB identifiers, state of the protein and whether it is bound to a ubiquitin variant, full length (FL), catalytic domain (CD) and cofactors and resolution.

|  | PDB-ID | Ubiquitin variant |  | Resolution |
| --- | --- | --- | --- | --- |
| **USP1** | 7AY2 | Ub<sup>Propargyl</sup> | CD + UAF1 | 3.2Å |
| **USP2** | 2HD5 | Ub<sup>wt</sup> | CD | 1.85Å |
| **USP4** | 2Y6E | Apo-form | CD (D1D2) | 2.4Å |
| **USP7** | 1NBF | Ub<sup>Aldehyde</sup> | CD | 2.3Å |
| **USP8** | 2GFO | Apo-form | CD | 2.0Å |
| **USP9X** | 5WCH | Apo-form | CD | 2.5Å |
| **USP12** | 5L8W | Ub<sup>Bromoethylamine</sup> | FL + UAF1 | 2.8Å |
| **USP14** | 2AYO | Ub<sup>Aldehyde</sup> | CD | 3.5Å |
| **USP15** | 7R2G | Active form | CD (D1D2) | 1.98Å |
| **USP18** | 5CHV | ISG15 | CD | 3.0Å |
| **USP21** | 2Y5B | Linear di-Ub<sup>Aldehyde</sup> | CD | 2.7Å |
| **USP25** | 6HEI | Ub<sup>Propargyl</sup> | CD | 1.64Å |
| **USP28** | 6HEK | Ub<sup>Propargyl</sup> | CD | 3.03Å |
| **USP30** | 5OHK | Ub<sup>Propargyl</sup> | CD | 2.34Å |
| **USP35** | 5TXK | Ub<sup>wt</sup> | CD | 1.84Å |
| **USP46** | 5L8H | Ub<sup>vinylmethylester</sup> | Full length | 1.85Å |

### Ub-Rhodamine activity assays

Enzyme activities were tested on a minimal substrate consisting of ubiquitin linked to a quenched fluorophore (Ub^Rho^, UbiQ). The USP40 activity was tested on Ub^AMC^ (UbiQ) instead of Ub^Rho^, as USP40 suffered from auto-inhibition when using Ub^Rho^ (Kim et al., in preparation). Enzyme activities were measured by the increase of fluorescence after cleavage (Rhodamine: Excitation at 485 nm, emission at 520 nm, AMC: Excitation at 350 nm, emission at 450 nm). All reactions were performed in a 384-well plate (Corning, flat bottom, low flange) on a Pherastar plate reader (BMG labtech) at room temperature. The assay buffer for USP1, USP7 and USP48 consisted of 20 mM Hepes, 150 mM NaCl, 5 mM DTT, 0.05% Tween-20. For USP15 and USP40 the assay buffer consisted of 20 mM Hepes pH 7.5, 100 mM NaCl, 5 mM DTT, 0.05% Tween-20.

For the kinetic analysis, we used defined enzyme concentrations of the different USPs related to intrinsic activity (USP7: 1 nM, USP1/UAF1, USP15^D1D2^ and USP40: 10 nM, USP48: 50 nM). Each USP variant was tested against a substrate concentration series generated by two-fold dilutions (USP1/UAF1 and USP40: 2500 nM to 39.1 nM, USP7: 1250 nM to 39.1 nM, USP15: 4000 nM to 62.5 nM, USP48: 3750nM to 234.4 nM). Substrate was prepared at 2x concentrations, after which 10µl of each substrate concentration in triplicates was pipetted to the 384 well-plate. Enzyme was injected to the plate at a 2x concentration using the Pherastar plate reader built in syringe. Measurement was started after the enzyme was injected to the plate. Durations of the different experiments varied, to ensure reactions ran to completion by fully hydrolyzing Ub^Rho^ or Ub^AMC^ (USP1/UAF1: 1890 sec, USP7: 3969 sec, USP15: 1764 sec, USP40 and USP48: 3600 sec).

### Michaelis-Menten analysis of USP activity

Fluorescence intensity data from the Ub^Rho^ activity assays were converted to substrate concentrations using a calibration curve. For each USP, calibration curves were generated using wildtype enzymes, where the plateau of the completed reactions resembles the concentration of released rhodamine. The converted data of the Ub^Rho^ activity assays were then analyzed by using the Michaelis-Menten model in GraphPad prism 7. First, initial velocities were determined from the linear phase of the reaction (first three datapoints). Then, in order to determine the kinetics, these initial velocities were plotted against the substrate concentration and then fitted using the non-linear regression Michaelis-Menten model of GraphPad Prism 7 software. In order to verify the results of the Michaelis-Menten analysis, we performed a global fit analysis of the data using Kintek Explorer version 8.0 (Kintek Corporation)(Johnson, 2009) (*Supplementary figure 3, Supplementary table 2*).

### pH analysis

Activity of each USP was tested at pH 7.0, 8.0 and 9.0, using assay buffer with 20 mM Hepes pH 7.0, 20 mM Hepes pH 8.0 or 20 mM MMT pH 9.0 replacing the 20 mM Hepes pH 7.5. Each USP was tested against a set concentration of Ub-Rho (1 µM). Enzymes and substrate were prepared at a 2x concentration in the three different buffers, and each reaction was performed in duplo. 10µl of enzyme in their different buffers were added to the plate. Measurement was started after 10µl of substrate was pipetted to the plate. A full kinetic analysis was performed on USP40^N452A^ and USP40^D453A^, (2500 nM to 39.06 nM) in assay buffer with 20 mM Hepes pH 8.0 to be compared with the earlier full kinetic analysis in assay buffer containing 20mM Hepes pH 7.5.

### PCNA-Ub deubiquitination assays

Activities of USP1/UAF1 and USP7 variants were tested against PCNA-Ub as a more complex substrate. Mono-ubiquitinated PCNA was produced with E1 and UbcH5C^S22R^ (UBE2D3) as described previously (Hibbert & Sixma, 2011) followed by purification on an S200 10/300 increase size exclusion chromatography column (GE healthcare). DUB activity assays were performed at room temperature with 2 µM PCNA-Ub in a reaction buffer composed of 20 mM Hepes pH 7.5, 150 mM NaCl, 2 mM DTT in a final volume of 210 µl. A 30µl sample was taken T=0 min sample before adding enzyme in order to get an accurate assessment of the ratio between PCNA-Ub and PCNA. In order to initiate the reaction, 100 nM of USP1/UAF1 or USP7 was added to remaining 180 µl reaction buffer containing 2 µM PCNA-Ub. Samples were taken at indicated time points and were added to SDS loading buffer in order to stop the reaction. Samples were loaded on a NuPAGE 4-12% Bis-Tris SDS gel and were separated by running them at 160V for 30 minutes. Gels were stained using Coomassie-Blue and were imaged using a Geldoc EZ imaging system (Bio-Rad Laboratories, Inc). Using Image Lab 6.0 software (Bio-Rad Laboratories, Inc), the volume intensities of non-ubiquitinated PCNA and ubiquitinated PCNA were measured for each time point. These volume intensities were then combined to calculate the total pool of PCNA in each lane, using which the percentage of non-ubiquitinated PCNA was determined. Since not all PCNA was ubiquitinated, all lanes were corrected for the percentage of non-ubiquitinated PCNA at t_0_.

### Expression of USP1 in RPE1 cells

RPE1 wildtype and USP1 knockout cells were a kind gift from Alan d’Andrea (Lim et al., 2018). USP1 knockout cells were lentivirally transduced with a doxycycline inducible USP1 expression vector (USP1^wt^, USP1^C90R^, USP1^D751A^ and USP1^D752A^). Cells were cultured in DMEM-F12. Transduced cells were selected with 10µg/ml blasticidin. Single clones with comparable USP1 expression to RPE1 wildtype cells were selected for each construct.

### Activity of USP1 mutants in RPE1 cells

Single clones for each construct were incubated with or without 1µg/ml doxycycline for 44 hours and were lysed using RIPA buffer (1% NP40, 1% sodium deoxycholate, 0.1% SDS, 0.15 M NaCl, 0.01 M sodium phosphate pH 7.5, 2 mM EDTA), containing cOmplete™, EDTA-free Protease Inhibitor Cocktail (Roche, 11873580001), 1mM 2-chloroacetamide and 0.25U/µl benzonase (SC-202391, Santa Cruz Biotechnology). Total protein concentration in the lysate was determined using a BCA assay (23227, Thermo Scientific) so that equal amounts could be loaded on gel. Samples were loaded on 4-12% Bolt gels (NW04127, Thermo Scientific), and run for 40 minutes at 180V in MOPS running buffer (B0001, Thermo Scientific). Proteins were transferred to nitrocellulose membrane (10600002, Amersham Protran 0.45 NC nitrocellulose). Membranes were stained with the following antibodies: Ubiquityl-PCNA (Lys164) (D5C7P) (13439, Cell Signaling Technology), PCNA (PC10) (sc-56, Santa Cruz Biotechnology), USP1 (14346-1-AP, Proteintech). After incubation with HRP coupled secondary antibodies the blots were imaged using a Bio-Rad Chemidoc XRS+. Using Bio-Rad ImageLab 5.1 software, we quantified PCNA-Ub levels by measuring the volume intensities of each PCNA-Ub band for each clone before and after doxycycline induction and ratios between these PCNA-Ub volume intensities were calculated. The distributions of these ratio values for different constructs were compared using non-parametric one-way ANOVA (Kruskal-Wallis test) corrected for multiple comparison by controlling the False Discovery Rate (Two-stage linear step-up procedure of Benjamini, Krieger and Yekutieli) in GraphPad Prism V9.5.

### Ubiquitin propargyl assays

Each USP variant (wt and mutants) was incubated at a single concentration (2 µM) with 16 µM of ubiquitin-propargyl (Ub^PA^, UbiQ) in crosslinking buffer (20 mM Tris pH 8.0, 150mM NaCl, 2 mM TCEP) at room temperature. Enzymes and substrate (Ub^PA^) were prepared at a 2x concentration in crosslinking buffer and were combined to initiate the reaction. As a reference, wildtype of each enzyme at 2x concentration was incubated with crosslinking buffer instead of Ub^PA^. After 5 minutes, samples were taken and added to SDS loading buffer to terminate the reaction. Samples were run on a 4-12% gradient gel.

## Results

### Positioning of catalytic residues in catalytic cleft is highly similar in different USPs

Directly adjacent to the canonical third catalytic residue (first critical residue), which was based on original findings in USP7 (Hu et al., 2002), lies a much better conserved aspartate (the second critical residue) (*Figure 1ABCD, Supplementary 1A*). This aspartate is present in all USPs except CYLD and USP50. USP50 does not harbor either of these two residues and is considered to be inactive.

**Figure 1:**
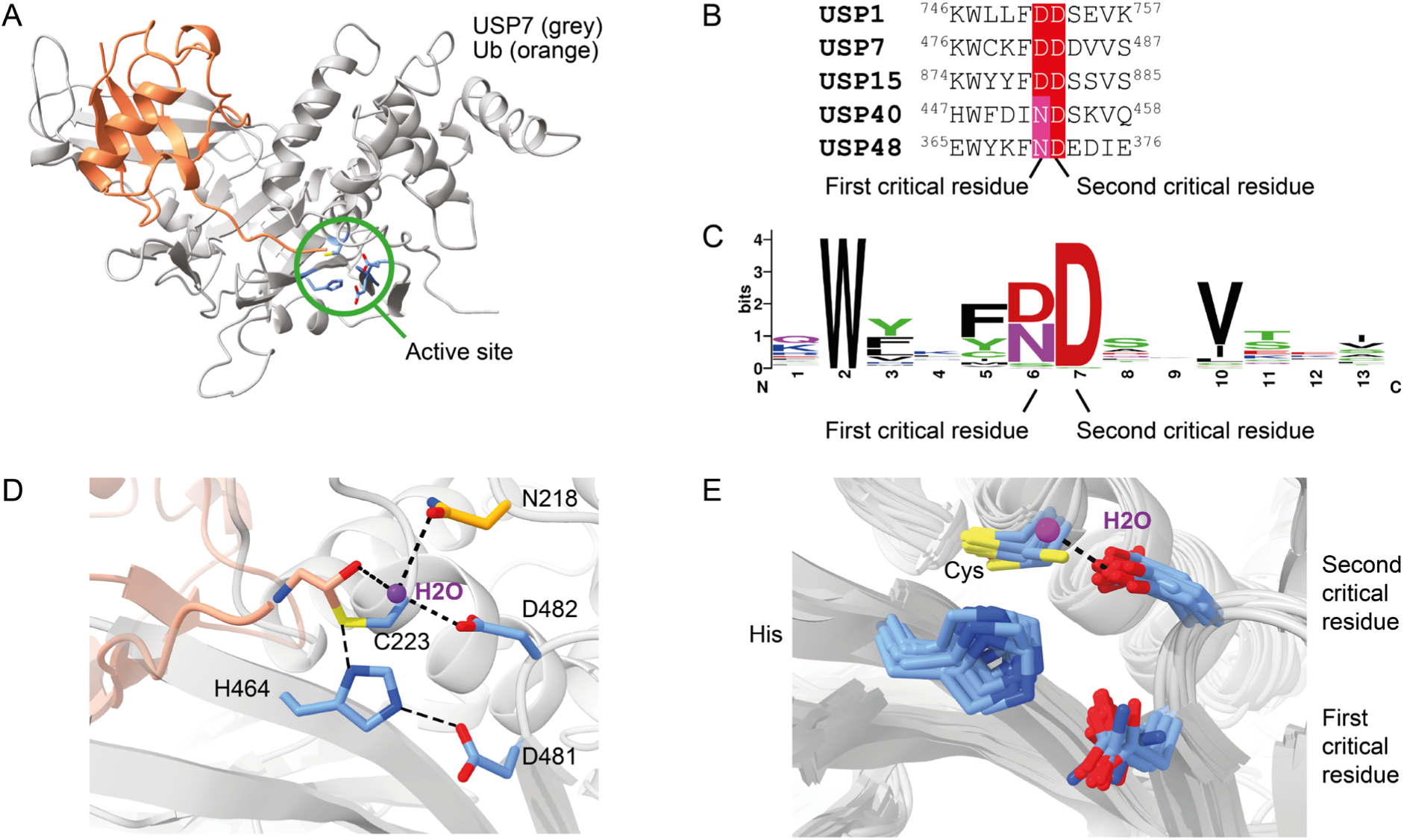
Structural and sequence alignment of USPs shows that both critical residues are positioned close to the catalytic histidine. **(A)**: Crystal structure of USP7 (grey) bound to ubiquitin-aldehyde (orange) (PDB: 1NBF). Location of the active site is highlighted (green) and catalytic residues, with the adjacent extremely conserved aspartate are shown in blue. **(B)**: Consensus of the catalytic mechanism shown by USP7 in complex with ubiquitin-aldehyde (1NBF, USP7: grey, ubiquitin-aldehyde: Pink). Catalytic cysteine, histidine, first critical residue and second critical residue are shown in blue. Hydrogen bonds (<3.5Å) are shown as black dashed lines. The oxygen atom of the first critical residue (D481) forms a hydrogen bond with the nitrogen (Nε) on catalytic histidine (H464), which allows histidine to activate cysteine for the nucleophilic attack. A water molecule (purple) is held in place by the second critical residue and an asparagine (N218, orange). This water molecule acts as member of the oxyanion hole to stabilize ubiquitin (pink). **(C)**: Sequence alignment of the five human USPs relevant for this study. **(D)**: Sequence logo of the 56-known human USPs, the canonical third catalytic residue (6) is referred to as first critical residue. The highly conserved adjacent residue (7) is referred to as second critical residue. Full sequence alignment is shown in Supplementary 1B. **(E)**: Superposition of available USP structures (Table 1) with aligned catalytic triads shows subtle variations in positioning of the residues. As the structure of USP35 has a cysteine to serine mutation, this residue is not shown. Hydrogen bonding of the second critical residue with water molecule is only present in USP7.

When structures of USP catalytic domains are superimposed (*Table 1, Figure 1E*) we can observe only minor differences in the positioning of these two adjacent residues. However, the interaction of the second critical residue with water, required for oxyanion hole formation, is exclusively found in USP7 despite the fact that some of these structures should have sufficient resolution to identify waters (Table 1).

Here we study the role of these adjacent critical residues in a selection of USPs. To allow accurate functional assessment in a side-by-side comparison, we generated the following mutations: USP1 (Res1: D751A, Res2: D752A), USP7 (Res1: D481A, Res2: D482A), USP40 (Res1: N452A, Res2: D453A), USP48 (Res1: N370A, D371A) and the D1D2 catalytic core of USP15 (Priyanka et al., 2022; Ward et al., 2018), USP15^D1D2^ (Res1: D879A, Res2: D880A).

As quality control we assessed potential misfolding or instability of all mutants tested using a thermal stability assay (*Supplementary figure 1B*). Comparing the individual critical residue loss of function mutants to their wildtype counterpart, we can observe only minor differences in thermal stability. While both mutants of USP15 have a decreased thermal stability compared to USP15^wt^, these variants retain stability to 50°C, indicating that they are still well-folded and suitable for kinetic assays at room temperature.

### The first critical residue is dispenUSP1/UAF1, USP15, USP40 and USP48 are active without the first critical residue

In order to measure the catalytic activities of wildtype and mutants of each USP, we followed enzyme activity using fluorogenic substrates, ubiquitin rhodamine (Ub^Rho^) or ubiquitin AMC (Ub^AMC^), at varying concentrations (*Figure 2, Supplementary figure 2*). We then studied the kinetics of these variants by Michaelis-Menten analysis (*Supplementary table 1*) and compared their catalytic efficiencies in order to study the importance of both critical residues (*Table 1*). Surprisingly, our experiments reveal that USP1/UAF1, USP15, USP40 and USP48 do not rely on the canonical third catalytic residue (first critical residue) and instead, are rendered catalytically dead only when their second critical residue is mutated.

**Figure 2:**
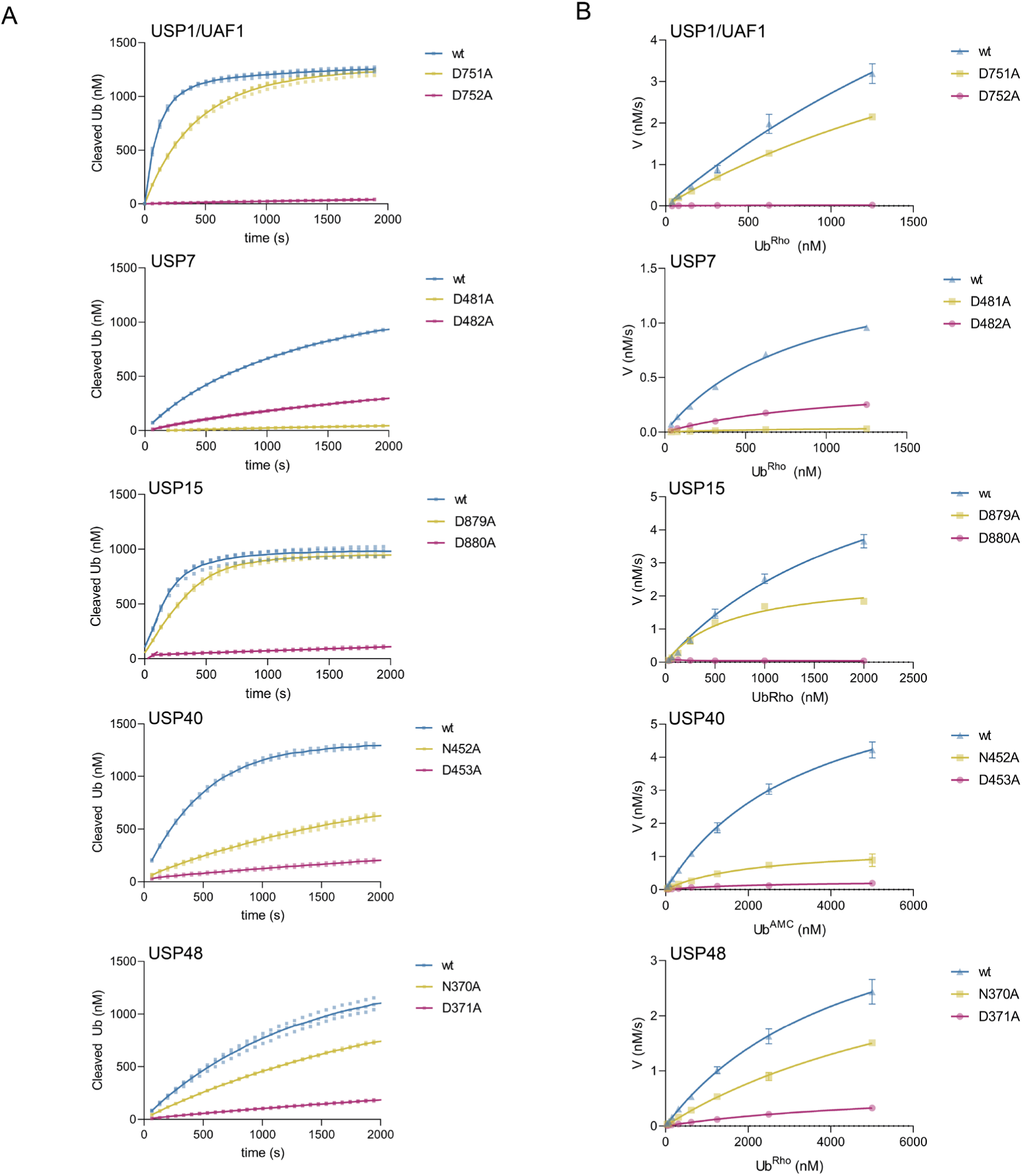
USP1, USP15, USP40 and USP48 do not rely on the canonical first critical residue. **(A)**: Enzyme activity assays of USPs on a minimal substrate (Ub^Rho^ for USP1/UAF1, USP7, USP15 and USP48, Ub^AMC^ for USP40). For each USP we show a single enzyme concentration against a single concentration of substrate. Wildtype (blue), mutation of first critical residue (yellow) and second critical residue are shown (purple). Assays with full range of substrate concentrations are shown in supplementary figure 2 (n=2, biological replicates, n=3, technical replicates). **(B)**: Michaelis-Menten analysis of enzyme activity assays shown in Supplementary 2.

Examination of the Michaelis-Menten analysis shows that USP1, USP15, USP40 and USP48 are all still catalytically competent upon loss of their first critical residue (*Figure 2, Table 2*). The loss of the first critical residue leads to minor decreases in their catalytic efficiency, which is slightly different for each USP. In USP15, we can observe that USP15^D879A^ does not decrease its catalytic efficiency at all compared to USP15^wt^. USP1^D751A^ only suffers from a minor 1.4-fold decrease in catalytic efficiency compared to USP1^wt^, and USP48^N370A^ suffers a 2-fold decrease compared to USP48^wt^. This effect is slightly bigger in USP40^N452A^, which decreases its catalytic efficiency 4-fold. Interestingly, out of the five USPs we tested, USP7 was the only USP where we observe a major effect on catalytic efficiency. Mutagenesis of the first critical residue (USP7^D481A^) leads to a large (26-fold) decrease in catalytic efficiency (kcat/KM) compared to USP7^wt^, rendering USP7^D481A^ mostly inactive, which agrees with existing literature on USP7 (Hu et al., 2002).

**Table 2:** Catalytic efficiencies (k_cat_/KM) of USPs and their critical residue mutants on a minimal substrate. Full Michaelis-Menten analysis is shown in supplementary table 1. Analogous Kintek verification of Michaelis-Menten analysis is shown in supplementary figure 2.

|  | Wildtype | Critical residue 1 | Critical residue 2 |
| --- | --- | --- | --- |
| | ($s^{-1}\mu M^{-1}$) | ($s^{-1}\mu M^{-1}$) | ($s^{-1}\mu M^{-1}$) |
| **USP1/UAF1** | 0.35 ($\pm 0.17$) | 0.24 ($\pm 0.013$) | 0.005 ($\pm 0.00036$) |
| **USP7** | 1.87 ($\pm 0.22$) | 0.07 ($\pm 0.008$) | 0.43 ($\pm 0.047$) |
| **USP15<sup>D1D2</sup>** | 0.35 ($\pm 0.074$) | 0.38 ($\pm 0.102$) | <0.0001 |
| **USP40** | 0.21 ($\pm 0.0065$) | 0.06 ($\pm 0.0073$) | 0.01 ($\pm 0.0035$) |
| **USP48** | 0.02 ($\pm 0.0009$) | 0.01 ($\pm 0.001$) | 0.002 ($\pm 0.0002$) |

### The second critical residue is essential for catalysis in USP1, USP15, USP40 and USP48

In contrast, our analysis reveals that the other USPs suffer a significant decrease in catalytic efficiency when their second critical residue is mutated, which results in USP1^D752A^, USP15^D880A^, USP40^D453A^ and USP48^D371A^ showing virtually no activity. Catalytic efficiency of USP1^D752A^ is 86-fold lower compared to USP1^wt^. In USP15^D880A^ we were not able to measure a signal of released fluorescent rhodamine at all, indicating that there is no activity left. Mutation of D453A in USP40, i.e. loss of the second critical residue, causes a 20-fold decrease in catalytic efficiency compared to USP40^wt^. USP48^D371A^ shows a 10-fold decrease in catalytic efficiency compared to USP48^wt^, which itself has a relatively low catalytic efficiency on Ub^Rho^, even while using a higher enzyme concentration (100 nM). Still, this 10-fold decrease renders USP48^D371A^ almost catalytically dead.

We find that the second critical residue, presumably its role in oxyanion intermediate stabilization, is only of minor importance in USP7, as seen by a minor (4-fold) decrease in catalytic efficiency of USP7^D482A^ compared to USP7^wt^, a smaller decrease than previously seen for USP7^D482A^ (Hu et al., 2002). These results do confirm that the first critical residue in USP7 makes up the catalytic triad. Our Michaelis-Menten analysis therefore implies that the role of the second critical residue in stabilization of the tetrahedral intermediate in USP7 is less important than was first thought.

Taken together, our findings demonstrate that for most USPs, the importance of this third catalytic residue is dispensable, and that there is variety in catalytic importance of the two critical residues between different USPs. Except for USP7, the second critical residue is essential in the majority of USPs tested here. However, multiple sequence alignment and structural alignment (*Figure 1BCE*) do not provide a clear indication about which of the two critical residues is more important. Because of this, use of these alignments to predict the importance and role of the critical residue is not sufficient.

### Activity on natural substrates confirms different catalytic triad composition

In order to validate that USP1/UAF1 can really function without the canonical first critical residue, we compared the activity of USP1/UAF1 and USP7 on a natural substrate. Deubiquitination activity was tested on mono-ubiquitinated PCNA (PCNA-Ub), a well-known substrate of USP1/UAF1 (Huang et al., 2006) and a potential substrate of USP7 (Kashiwaba et al., 2015). We found that USP1/UAF1, USP7 and their mutants display the same relative activity towards PCNA-Ub as was observed for a minimal substrate. As expected, USP1^wt^/UAF1 and USP7^wt^ are able to cleave PCNA-Ub (*Figure 3AB*). The first critical residue mutant USP1^D751A^/UAF1 causes a minor decrease in cleavage of PCNA-Ub compared to USP1^wt^/UAF1. We find that just like their activity on Ub^Rho^, USP1^D752A^ (second critical residue) and USP7^D481A^ (first critical residue) are unable to cleave PCNA-Ub, with USP7^D481A^ still retaining minimal activity. Mutating the second critical residue in USP7^D482A^ displays only a minor decrease in activity compared to USP7^wt^, although this could be due to a difference in enzyme/substrate ratio. Taken together, the different relative importance of the first and second residue is retained on different substrates.

**Figure 3:**
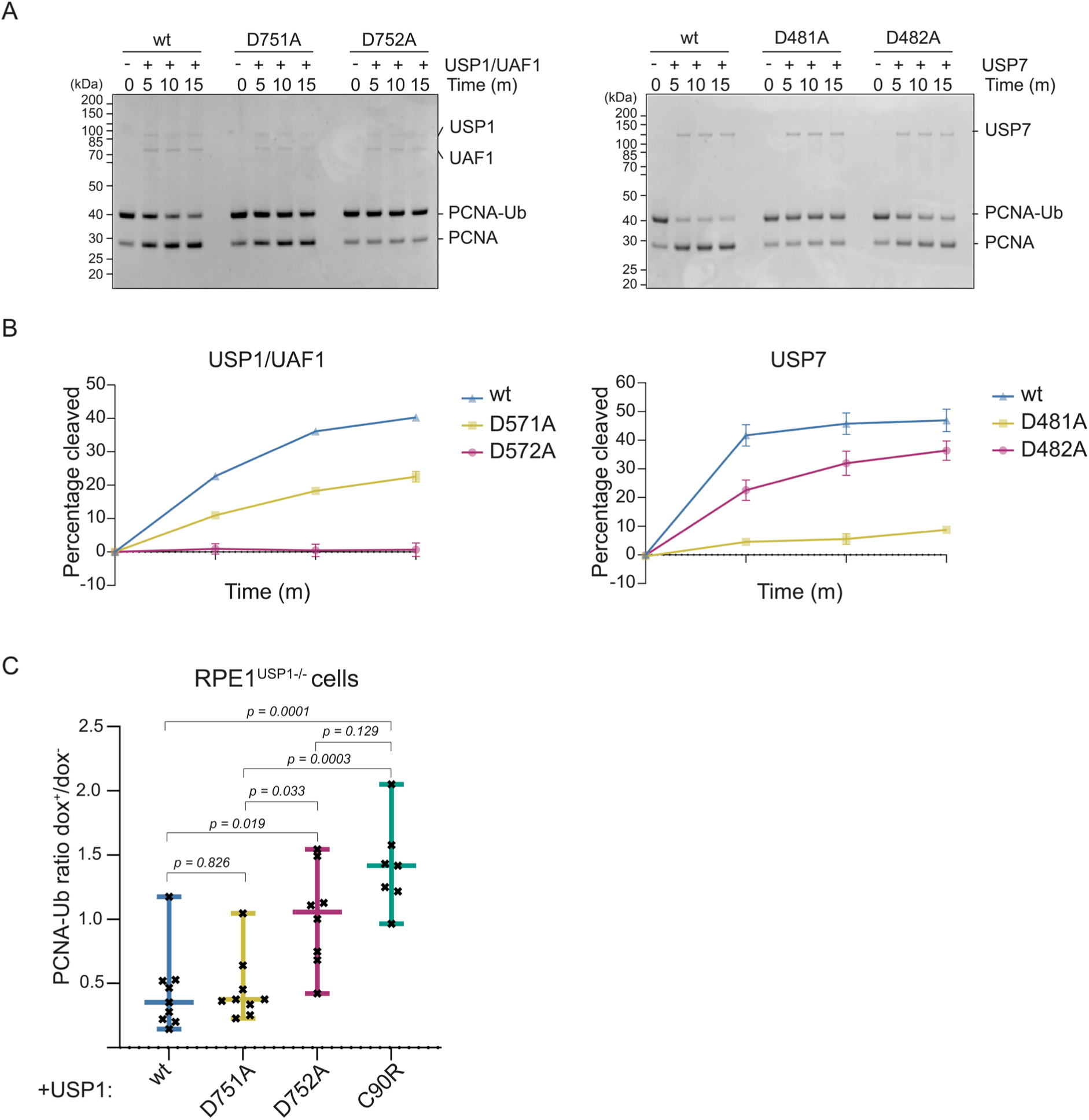
The varying use of critical residues in USP1 and USP7 is retained on a natural substrate and confirmed for USP1 in cells. **(A)**: In vitro deubiquitination assay of USP1/UAF1 and USP7 on a natural substrate (PCNA-Ub) comparing the importance of both critical residues. Results confirm that the first critical residue is more critical for USP7, and the second critical residue is more critical for USP1/UAF1. **(B):** Quantification of gel-based activity assays. USP1^D752A^ and USP7^D481A^ are catalytically incompetent, whereas USP1^D751A^ and USP7^D482A^ are still able to cleave PCNA-Ub. **(C):** Quantification of PCNA deubiquitination in USP1 RPE1 knockout cell lines complemented with USP1^wt^ and USP1^D751A^, USP1^D752A^ and USP1^C90R^ (Supplementary figure 4). Cell lysates were stained using antibodies for PCNA-Ub, PCNA and USP1. Levels of PCNA-Ub were quantified before (dox^-^) and after (dox^+^) doxycycline induction to determine the ratio of PCNA-Ub. P-values (see methods) are shown and results confirm that USP1^D751A^ behaves like wildtype and that USP1^D752A^ significantly loses activity.

### USP1 requires the second critical residue to process PCNA-Ub in cells

We wondered whether the importance of the second critical residue for catalysis in USP1 also holds up in a cellular context. Using doxycycline inducible lentiviral expression in mammalian (RPE1) cells, we complemented a USP1 knockout cell line with full-length USP1^wt^, USP1^D751A^ and USP1^D752A^ to investigate whether their activity is comparable to that seen on Ub^Rho^ and to *in vitro* activity on PCNA-Ub. As a control, cells were complemented with catalytically dead USP1^C90R^ to validate our experimental setup. Between single clones of our USP1 knockout cell line, we observed variable levels of basal PCNA-Ub even before inducing expression with doxycycline (*Supplementary figure 4*). We therefore calculated the ratio of PCNA-Ub levels before and after doxycycline induction within single clones, and provide ratios averaged over multiple clones. Expressing USP1^wt^ caused a decrease in PCNA-Ub levels whereas USP1^C90R^ expression did not affect PCNA ubiquitination levels, confirming that our experimental setup was functioning (*Figure 3C*).

Introducing USP1^D751A^ and USP1^D752A^ in the USP1 knockout cells confirms that even in cells, mutation of the first critical residue (USP1^D751A^) does not affect its catalytic activity. Instead, the second critical residue mutant (USP1^D752A^) loses almost all its activity, comparable to USP^C90R^. Taken together, these experiments validate the importance of the second critical residue for cellular substrate processing and suggest that the outcome of in vitro analyses of USP catalytic mechanisms translate into relevant cellular substrate turnover.

### Different catalytic triad compositions are not affected by pH

Next, we decided to assess the differential importance of both residues in more detail to define their precise roles during substrate processing. The ionization properties of catalytic residues, described by their pKa, are of vital importance for accommodating catalysis. We examined the effect of buffer pH on the activity of different mutants, since at elevated pH the catalytic cysteine is expected to be more deprotonated. This in turn would make the first step of the catalytic cycle less dependent on the other two residues of the catalytic triad. We performed the DUB activity assays at pH 7.0, pH 8.0 and pH 9.0 against a single concentration of minimal substrate (*Supplementary Figure 5A*). Results show that the relative importance of the critical residues in USP1, USP7, USP15 and USP48 is not influenced by a change in pH. Although these USPs are more active at pH 8.0 and pH 9.0, their diverging preference for the different critical residue remains.

In USP40 the mutants appear to be more pH sensitive than USP40^wt^, with USP40^N452A^ and USP40^D453A^ having lower activity at pH 7.0 and pH 9.0, but a higher activity at pH 8.0. Additionally, at a higher pH (8.0 and 9.0), activity of USP40^N452A^ and USP40^D453A^ become more similar while at a lower pH (7.0) USP40^N452A^ retains more activity than USP40^D453A^, and this is also seen when comparing pH 8.0 to their activity to the previously tested pH 7.5 (*Supplementary Figure 5B*). These results suggest that the second critical residue in USP1, USP15, USP40 and USP48 and the first critical residue in USP7 are involved in cysteine deprotonation regardless of pH.

### Both critical residues promote polarization of the catalytic histidine in USP7, USP15 and USP40

Cleaving a ubiquitin-substrate bond is not a single event, but instead a series of events which culminates in the release of ubiquitin and the substrate. The two adjacent critical residues play different roles in this process, either polarizing histidine, which in turn allows cysteine to attack the ubiquitin-substrate linkage, or resolving the tetrahedral intermediates formed after the nucleophilic attack. We used ubiquitin-propargyl (Ub^PA^) to directly analyze the first step, the nucleophilic attack and eliminate the need to resolve the tetrahedral intermediates to complete the catalytic cycle. In doing so, we were able to specifically investigate which critical residue is necessary to polarize the catalytic histidine. We compared the ability of the different USP variants to crosslink to Ub^PA^ (*Figure 4*).

**Figure 4:**
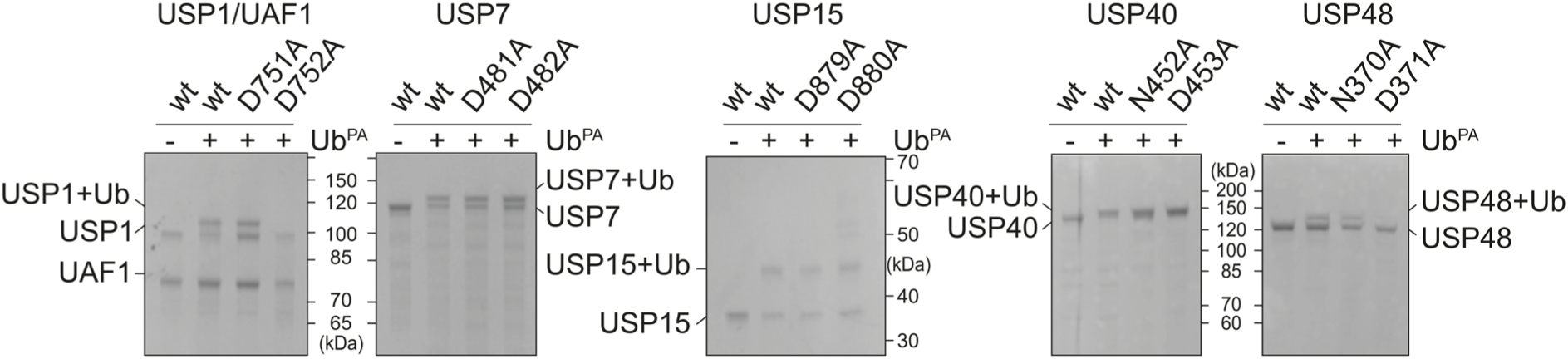
Both critical residues can perform nucleophilic attack in USP7, USP15 and USP40 and only the second residue can do this in USP1 and USP48. Analysis of ability of USPs and their critical residue mutants to successfully crosslink to ubiquitin-propargyl (Ub^PA^). Only the second critical residue is able to accommodate crosslinking to Ub^PA^ in USP1/UAF1 and USP48, but both critical residues can accommodate crosslinking in USP7, USP15 and USP40.

Interestingly, the catalytic cysteine in USP1 and USP48 is unable to crosslink when the second critical residue is lacking (USP1^D752A^ and USP48^D371A^), but can still crosslink when the first critical residue is mutated (USP1^D751A^ and USP48^N370A^). This is noteworthy as the propargylamine warhead does not feature the carbonyl of Gly76 which would be stabilized by the oxyanion hole residues. This suggests that the second critical residue can polarize the catalytic histidine and thereby accommodate the nucleophilic attack. In contrast, all USP7, USP15 and USP40 variants are able to crosslink to Ub^PA^. This implies, that either critical residue is able to promote polarization of the histidine and activate the catalytic cysteine for crosslinking. In summary, these data demonstrate that the catalytic mechanism employed by different members of the USP family is remarkably variable.

## Discussion

In this study, we investigated the catalytic mechanism employed by USP DUBs and show how the two critical residues contribute to catalysis with surprising variety. By combining data from the crosslinking experiment (*Figure 4*), which shows which critical residue is involved in the nucleophilic attack, with the activity assay on Ub^Rho^ (*Figure 2*), which shows which residue is required for performing full catalysis, we can make a model of the contribution of these two residues to catalysis (*Figure 5*).

**Figure 5:**
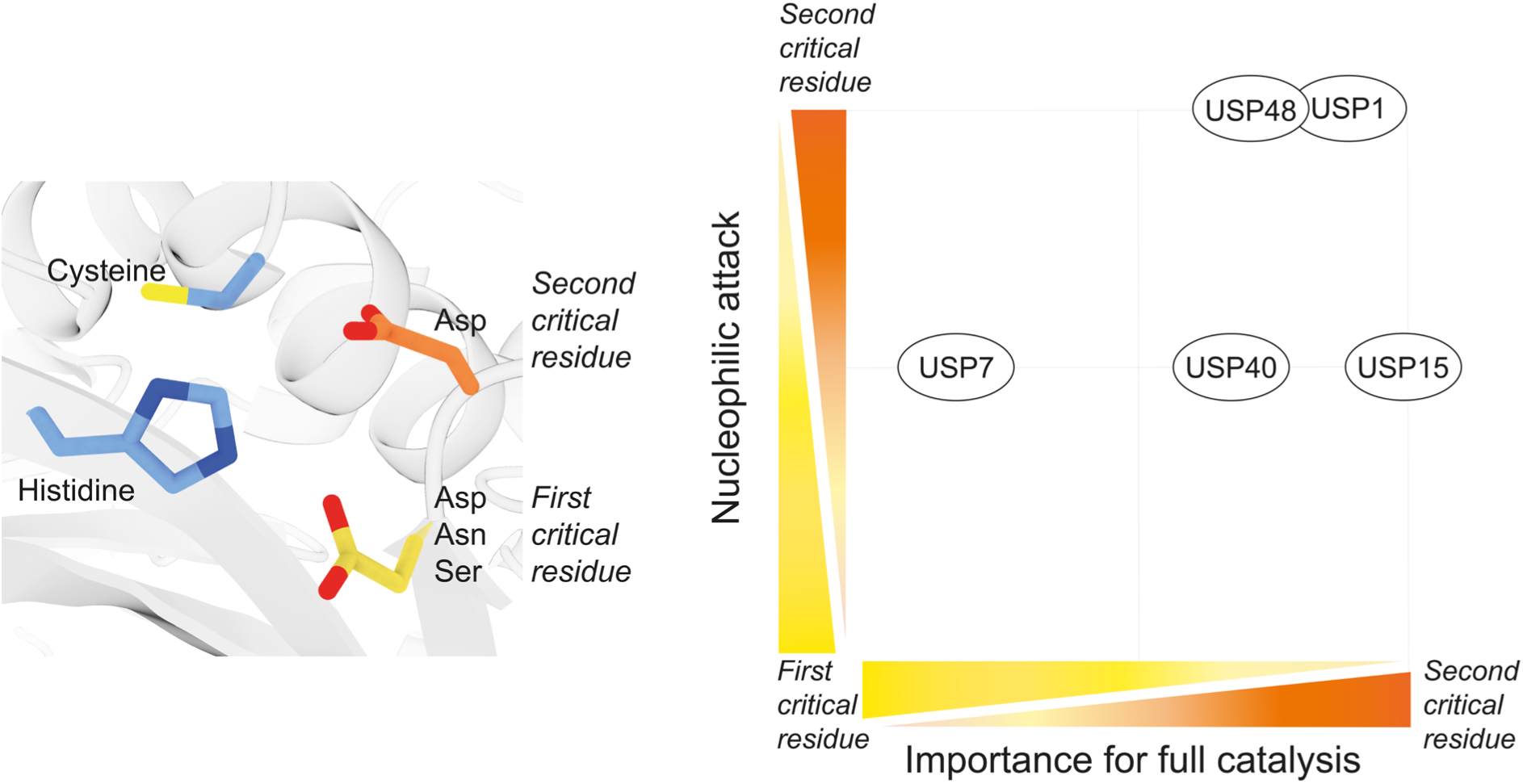
The critical residues perform different functions in different USPs. Depiction of the catalytic residues. Catalytic cysteine and histidine are shown in blue. The first and second critical residue are shown in yellow and orange respectively. Ability of the critical residue to accommodate the nucleophilic attack is shown on Y-axis, and importance of the critical residue for full catalysis is shown on the X-axis. In USP1 and USP48, only the second critical residue is able to polarize the catalytic histidine leading to a successful nucleophilic attack. Both critical residues are able to polarize the catalytic histidine in USP7, USP15 and USP40. After the nucleophilic attack, several tetrahedral intermediates need to be resolved in order to complete the catalytic cycle. In USP15, the second critical residue is required to complete the catalytic cycle on its own, although both residues are able to polarize histidine in USP15. USP1, USP40 and USP48 rely mostly on the second critical residue, but require some varying involvement of the first critical residue as well. USP7 primarily relies on the first critical residue and requires a minor contribution of the second critical residue.

The model requires two axes, denoting relative importance for nucleophilic attack (y-axis) and full catalysis (x-axis). In USP1 and USP48, we can observe that only the second critical residue is able to accommodate the nucleophilic attack, which is in line with its importance for full catalysis. In USP7, USP15 and USP40, both critical residues are able to accommodate the nucleophilic attack on their own (y-axis). Ability to polarize does not guarantee catalytic competence, as USP15 and USP40 rely mostly on the second critical residue when cleaving Ub^Rho^ (x-axis). Only USP7 lies on the other end of the spectrum, relying on the first critical residue for full catalysis and requires only a minor contribution of the second critical residue, previously shown to be due to involvement in oxyanion stabilization (Hu et al., 2002). In short, these five USPs all make different use of the two critical residues for catalysis.

All in all, while this is only a small subset of USPs, our data demonstrates that the consensus USP catalytic mechanism, based on findings in USP7 (Hu et al., 2002), is not valid for the other USPs tested here and that there is a remarkable variability of catalytic mechanisms among USPs.

The majority of USPs tested can complete the entire catalytic cycle without the canonical third catalytic residue, indicating that these USPs are still able to deprotonate the catalytic cysteine to allow for the attack on the ubiquitin tail. USP7, USP15 and USP40 all three have misaligned catalytic triads (Faesen et al., 2011; Priyanka et al., 2022; Ward et al., 2018; Kim et al., in preparation), which suggests an intrinsic flexibility in the active site. This flexibility could allow both critical residues to position themselves for hydrogen bonding to the catalytic histidine, even though such an arrangement for the second critical residue has not been observed in any USP crystal structure and considering the relative arrangement would be difficult to foresee without structural plasticity. Alternatively, hydrolase activity by a Cys-His dyad as is commonly observed in various other proteases could be sufficient in the context of strong substrate activation by the oxyanion hole’s second critical residue. There might be other USPs which have a misaligned catalytic triad, or in which the active site allows for more ’breathing’ than in other USPs. Such a flexibility could be missing in USP1 and USP48, where only the second critical residue can accommodate the nucleophilic attack.

However, the ability of USP7^D481A^, USP15^D880A^ and USP40^D453A^ to accommodate the nucleophilic attack does not guarantee catalytic competence, as shown by their lack of activity on Ub^Rho^. It is possible that after these mutants accommodate the nucleophilic attack, they would get stuck trying to resolve the tetrahedral intermediates. Thus, after a single nucleophilic attack, they would be unable to release ubiquitin and the substrate and would therefore be unable to proceed to cleave the next substrate.

The main difference between the critical residue mutants in these USPs is then their ability to complete the catalytic cycle, for which the Ub^Rho^ activity assays offer a more definitive insight into the importance of the critical residues. Of note, while fluorescence is generated in parallel to generation of the thioester intermediate, the signal observed in our assay requires multi-turnover conditions and thus corresponds to the full catalytic cycle.

The presence of a negative charge in the first critical residue does not affect its importance for catalysis. Our analysis shows that of the USPs with an aspartate as their first critical residue (USP1, USP7 and USP15), only USP7 depends on it for catalysis. USP40 and USP48 have an asparagine as their first critical residue, and this asparagine does not appear to be crucial for catalysis. While most USPs harbor an asparate or asparagine, three USPs, USP16, USP30 and USP45, harbor a serine in position of the first critical residue instead and are all three are catalytically competent (Joo et al., 2007; Gersch et al., 2017; Perez-Oliva et al., 2015). The role of serine itself was only studied in USP30 and when serine was substituted for asparagine and especially for aspartate, there was a major effect on catalysis and a dampening of its K6-linkage selectivity (Gersch et al., 2017). As serine is highly efficient at forming H-bonds, it could be that this serine is the critical residue of choice in USP16 and USP45 as well. Recently, cellular USP45 was shown to display elevated reactivity towards protein probes, not in line with its in vitro catalytic activity, which stresses that serine as the third catalytic residue does not coincide with lower nucleophilicity (O’Dea et al., 2023).

The other role of the second critical residue is formation of the oxyanion hole. A dual role, with a single critical residue stabilizing catalytic histidine and oxyanion hole formation simultaneously is unlikely, as histidine stabilization by the critical residue is required throughout the entire catalytic cycle. Interestingly, our data shows that the majority of tested USPs are able to perform catalysis using just the second critical residue, and that this is not possible with just the first critical residue. It is possible that some USPs have different requirements for oxyanion stabilization, due to minor differences in residues surrounding the catalytic cleft. USPs have an extremely conserved asparagine (USP2: N271, PDB: 2HD5, USP7: N218, PDB: 1NBF) structurally poised to stabilize oxyanion intermediates (Hu et al., 2002; Zhang et al., 2011). This asparagine alone could be sufficient for stabilizing the oxyanion intermediate in some USPs.

Direct hydrogen bonding of the second critical residue with histidine would not allow for catalysis, as this requires the histidine to flip, which would leave no nitrogen positioned for cysteine deprotonation. This was also shown in earlier research, where disruption of the active site by an allosteric inhibitor causes the histidine to flip (Rennie et al., 2022), resulting in inactive USP1. Hydrogen bonding histidine would have to take place via an alternative water-mediated interaction. Through this water molecule, the second critical residue could act as a base, and would be able to reach catalytic histidine or even the catalytic cysteine. Interestingly, this second critical residue is almost always an aspartate. The high sequence and structural conservation of this second critical residue (aspartate) implies an importance of its negative charge. This negative charge itself could contributes to the catalytic mechanism and promote protonation and deprotonation of the other residues involved.

Our findings are important for fundamental understanding of USP DUB function. This paper highlights the importance of mutagenesis in order to characterize the catalytic triad as structural analysis alone does not explain the catalytic mechanism. Without appropriate characterization of the catalytic triad, research could be focused on the wrong residues. Additionally, inaccurate assumptions about the catalytic triad in USP could affect conclusions made regarding loss of function mutations in genetic screens.

The observed variability of mechanisms in USPs may also open up new opportunities for drug discovery. Targeting USPs is a viable strategy for treatment of many different diseases, with promising USP inhibitors currently under development (Cadzow et al., 2020; Tsefou et al., 2021). In addition, the possibilities for DUBTACs (Henning et al., 2022), make these exploits even more relevant. The remarkable variability of catalytic mechanisms may make it possible to selectively target individual USPs.

## Acknowledgements

The authors would like to thank Farid El Oualid, other members of the department and Herbert Waldmann for all the useful discussions. Funding was provide by NWO (OCENW.KLEIN.131 and LIFT 731.017.415) and Oncode Institute.

