## supplemental data for "Surprising variety in the USP deubiquitinase catalytic mechanism"

### Supplementary data

***Supplementary table 1: Analysis of kinetics of USPs and their mutants using the Michaelis-Menten equation.*** *Comparisons between different USPs and their mutants should be made by comparing $k_{cat}/K_M$, not individual $k_{cat}$ and $K_M$ as in this experimental set up, $k_{cat}$ affects $K_M$.*

| | | **$k_{cat}$ ($s^{-1}$)** | **$K_M$ (µM)** | **$k_{cat}/K_M$** |
|---|---|---|---|---|
| **USP1** | **wt** | 1.27 (±0.38) | 3.68 (±1.40) | 0.345 (±0.17) |
| | **D751A** | 1.47 (±0.03) | 5.36 (±0.15) | 0.245 (±0.01) |
| | **D752A** | 0.0036 (±0.00008) | 0.54 (±0.034) | 0.005 (±0.0004) |
| **USP7** | **wt** | 1.65 (±-0.093) | 0.88 (±0.092) | 1.873 (±0.22) |
| | **D481A** | 0.047 (±0.002) | 0.69 (±0.073) | 0.069 (±0.008) |
| | **D482A** | 0.48 (±0.026) | 1.11 (±0.105) | 0.429 (±0.047) |
| **USP15** | **wt** | 0.80 (±0.09) | 2.31 (±0.42) | 0.346 (±0.074) |
| | **D879A** | 0.26 (±0.027) | 0.68 (±0.17) | 0.383 (±0.103) |
| | **D880A** | - | - | - |
| **USP40** | **wt** | 0.72 (±0.011) | 3.47 (±0.097) | 0.207 (±0.0065) |
| | **N452A** | 0.13 (±0.006) | 2.12 (±0.23) | 0.061 (±0.0073) |
| | **D453A** | 0.027 (±0.003) | 2.27 (±0.60) | 0.012 (±0.0035) |
| **USP48** | **wt** | 0.094 (±0.02) | 4.68 (±0.18) | 0.020 (±0.0009) |
| | **N370A** | 0.080 (±0.02) | 8.27 (±1.94) | 0.010 (±0.001) |
| | **D371A** | 0.17 (±0.014) | 6.22 (±0.747) | 0.002 (±0.0002) |

***Supplementary table 2: Verification of Michaelis-Menten analysis by using a global fit analysis using Kintek Explorer version 8.0*** *(Kintek Corporation* (Johnson et al., 2009). *Fluorescence intensity data of all the DUBs was fitted with the model that consists of three steps: substrate binding, and the combination of substrate cleavage and release of the label, and dissociation from the target Ub. Cleavage data from wildtype and all the mutants of each DUB was simultaneously fitted keeping the dissociation rates of the substrate ($Ub^{Rho}$ or $Ub^{AMC}$) the same for all the DUB variants also dissociation rates of the product (Ub) were kept the same. Concentrations substrate were allowed to vary up to 5% from their theoretical values in order to compensate for experimental pipetting error. The output of the models was verified by comparing values of the coefficients that were used by the model in order to convert fluorescence intensity units to product concentration units with the values of these coefficients that were calculated from calibration curves (M&M). The $K_M$, $k_{cat}$ and $k_{cat}/K_M$ values were derived from the reaction rate constants of individual reaction steps:*

$$k_{cat} = \frac{k_2 k_3}{k_2 + k_{-2} + k_3}$$

$$K_M = \frac{k_2 k_3 + k_{-1}(k_{-2} + k_3)}{k_1(k_2 + k_{-2} + k_3)}$$

| | $k_1$ (1/(μM*s))[a] | $k_{-1}$ (1/s)[c] | $k_2$ (1/s) | $k_{-2}$ (1/s)[b] | $k_3$ (1/s)[c] | $k_{-3}$ (1/(μM*s))[a] | $k_{cat}$ (1/s) | $K_M$ (μM) | $k_{cat}/K_M$ (1/(μM*s)) |
|---|---|---|---|---|---|---|---|---|---|
| **USP1 wt** | 100 | 105.0<br>± 6.8 | 2.080<br>± 0.071 | $10^{-10}$ | 40.8<br>± 4.1 | 100 | 1.979<br>± 0.210 | 1.03<br>± 0.13 | 1.942<br>± 0.316 |
| **USP1 D571A** | 100 | 105.0<br>± 6.8 | 0.989<br>± 0.065 | $10^{-10}$ | 40.8<br>± 4.1 | 100 | 0.966<br>± 0.116 | 1.03<br>± 0.14 | 0.933<br>± 0.170 |
| **USP1 D572A** | 100 | 105.0<br>± 6.8 | 0.004<br>± 0.0002 | $10^{-10}$ | 40.8<br>± 4.1 | 100 | 0.004<br>± 0.0005 | 1.05<br>± 0.14 | 0.004<br>± 0.001 |
| **USP7 wt** | 100 | 106.3<br>± 9.7 | 2.280<br>± 0.120 | $10^{-10}$ | 148.6<br>± 23.1 | 100 | 2.246<br>± 0.368 | 1.07<br>± 0.20 | 2.100<br>± 0.524 |
| **USP7 D481A** | 100 | 106.3<br>± 9.7 | 0.100<br>± 0.010 | $10^{-10}$ | 148.6<br>± 23.1 | 100 | 0.010<br>± 0.018 | 1.06<br>± 0.22 | 0.094<br>± 0.026 |
| **USP7 D482A** | 100 | 106.3<br>± 9.7 | 0.630<br>± 0.050 | $10^{-10}$ | 148.6<br>± 23.1 | 100 | 0.627<br>± 0.109 | 1.06<br>± 0.21 | 0.590<br>± 0.155 |
| **USP15 wt** | 100 | 100.0<br>± 11.6 | 0.181<br>± 0.012 | $10^{-10}$ | 100.0<br>± 11.6 | 100 | 0.181<br>± 0.024 | 1.00<br>± 0.18 | 0.181<br>± 0.04 |
| **USP15 D879A** | 100 | 100.0<br>± 11.6 | 0.118<br>± 0.008 | $10^{-10}$ | 100.0<br>± 11.6 | 100 | 0.118<br>± 0.016 | 1.00<br>± 0.18 | 0.118<br>± 0.026 |
| **USP40 wt** | 100 | 171.0<br>± 5.01 | 0.647<br>± 0.018 | $10^{-10}$ | 171.0<br>± 5.01 | 100 | 0.645<br>± 0.026 | 1.71<br>± 0.09 | 0.377<br>± 0.024 |
| **USP40 N452A** | 100 | 171.0<br>± 5.01 | 0.116<br>± 0.003 | $10^{-10}$ | 171.0<br>± 5.01 | 100 | 0.116<br>± 0.004 | 1.71<br>± 0.08 | 0.068<br>± 0.004 |
| **USP40 D453A** | 100 | 171.0<br>± 5.01 | 0.034<br>± 0.001 | $10^{-10}$ | 171.0<br>± 5.01 | 100 | 0.0341<br>± 0.001 | 1.71<br>± 0.09 | 0.018<br>± 0.001 |
| **USP48 wt** | 100 | 374.0<br>± 3.12 | 0.070<br>± 0.001 | $10^{-10}$ | 354.0<br>± 17.2 | 100 | 0.070<br>± 0.003 | 3.73<br>± 0.19 | 0.019<br>± 0.001 |
| **USP48 N370A** | 100 | 374.0<br>± 3.12 | 0.042<br>± 0.0003 | $10^{-10}$ | 354.0<br>± 17.2 | 100 | 0.042<br>± 0.002 | 3.74<br>± 0.19 | 0.011<br>± 0.001 |
| **USP48 D371A** | 100 | 374.0<br>± 3.12 | 0.007<br>± 0.0001 | $10^{-10}$ | 354.0<br>± 17.2 | 100 | 0.007<br>± 0.0003 | 3.74<br>± 0.19 | 0.002<br>± 0.0001 |

[a] values of association rate constants of substrate and product ($k_1$ and $k_{-3}$) were fixed to approximate diffusion limit

[b] Reverse reaction rate constants were fixed close to 0

[c] for each DUB one dissociation rate constant of the substrate was used to fit all the data for the wildtype and the mutant proteins. Same was done for dissociation of the product.

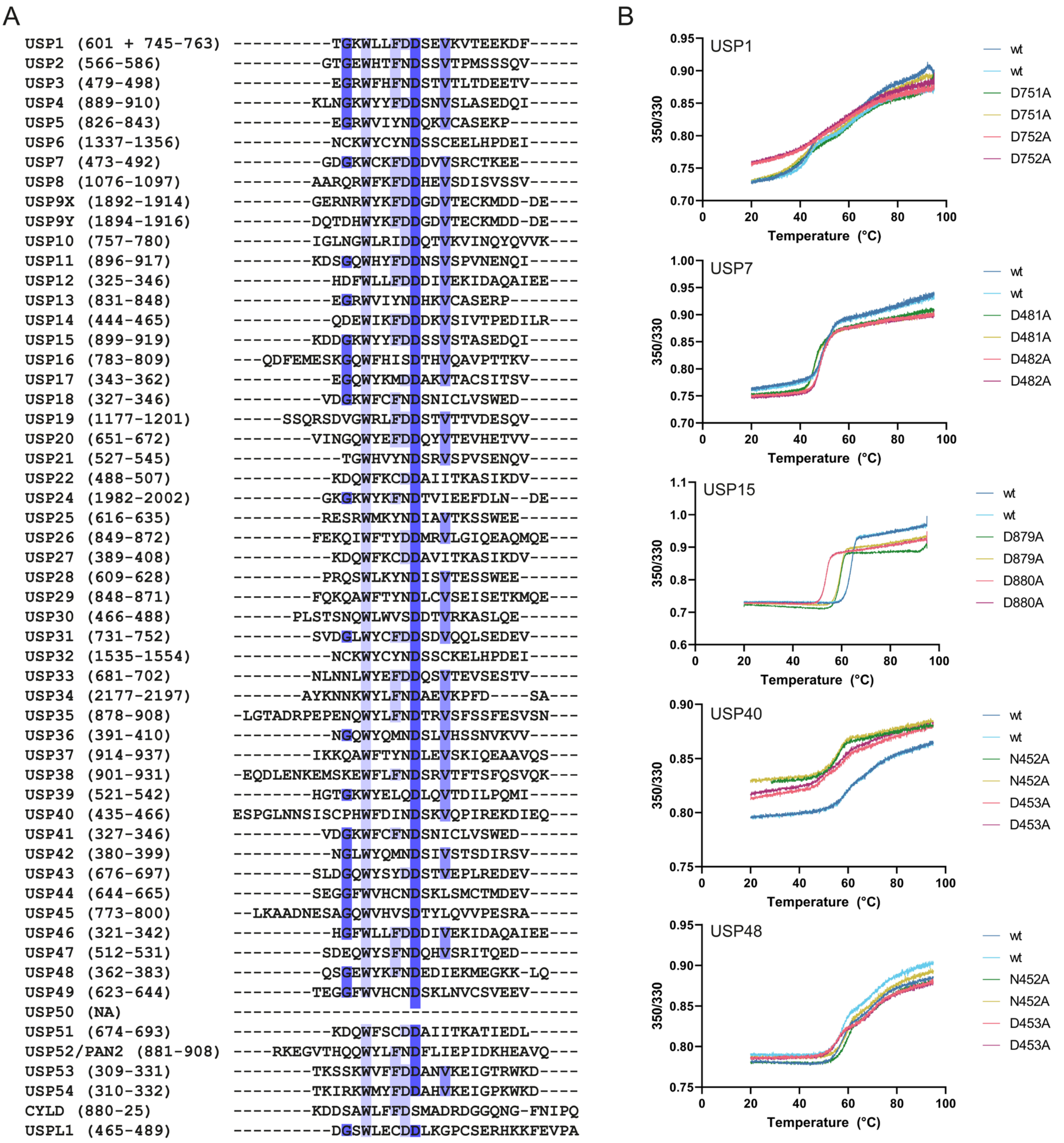

***Supplementary figure 1:*** ***(A):*** *Sequence alignment of all USP catalytic domains found on the UniProt database. Sequences are coloured based on percentage sequence Identity. The second critical residue is a highly conserved aspartate with over 96.4% sequence identity. Percentage identity for the first critical residue is lower, with 46.4% of the residues being an aspartate. 44.6% of USPs harbor an asparagine and 5.4% of USPs have a serine. USP39 is the only USP with a glutamate and is considered to be inactive* (Van Leuken et al., 2008)*. USP50 does not harbor the two adjacent residues and is therefore thought to be inactive* (Quesada et al., 2004)*.* ***(B):*** *NanoDSF thermal stability assay on all USPs and their mutants. USP1/UAF1 was tested as a heterodimer. Ratios measured (350mm/330mm) varied between some of the mutants. All proteins used in this study are stable above 40°C or higher and thus were stable under experimental conditions (25°C)*

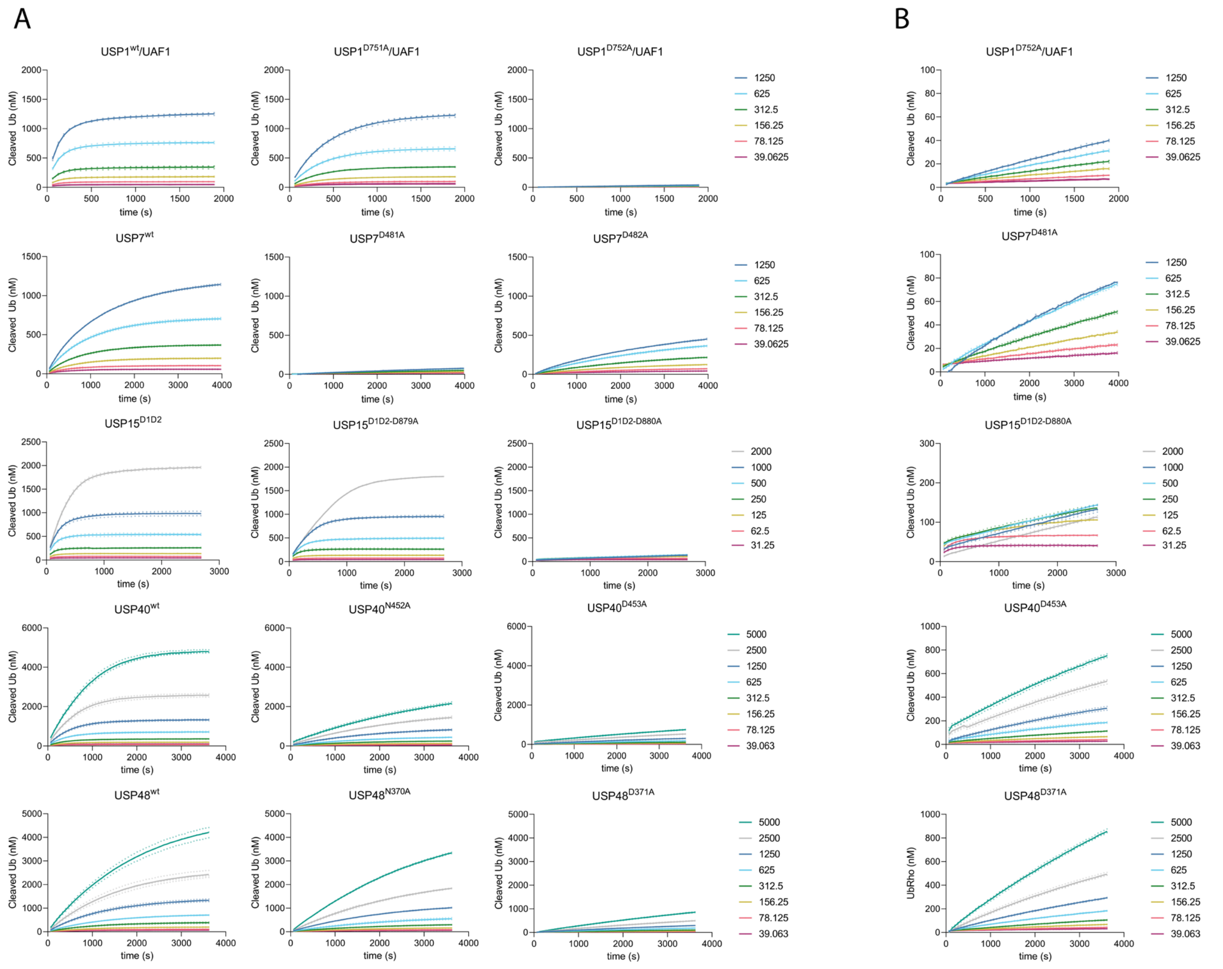

***Supplementary figure 2: Experimental data of enzyme activity assays performed on USP1/UAF1, USP7, USP15$^{D1D2}$, USP40 and USP48 and their mutants, used for Michaelis-Menten analysis.*** *Single enzyme concentrations were tested against a range of substrate concentrations as shown. Equivalent concentrations are given in identical colours. Reactions were run for varying durations to ensure each reaction ran to completion, which was needed in order to generate calibration curves as described in methods. Individual points were experimentally aquired, connecting lines are introduced for visualization purposes.* ***(B):*** *Zoom of the catalytically incompetent mutants.*

A

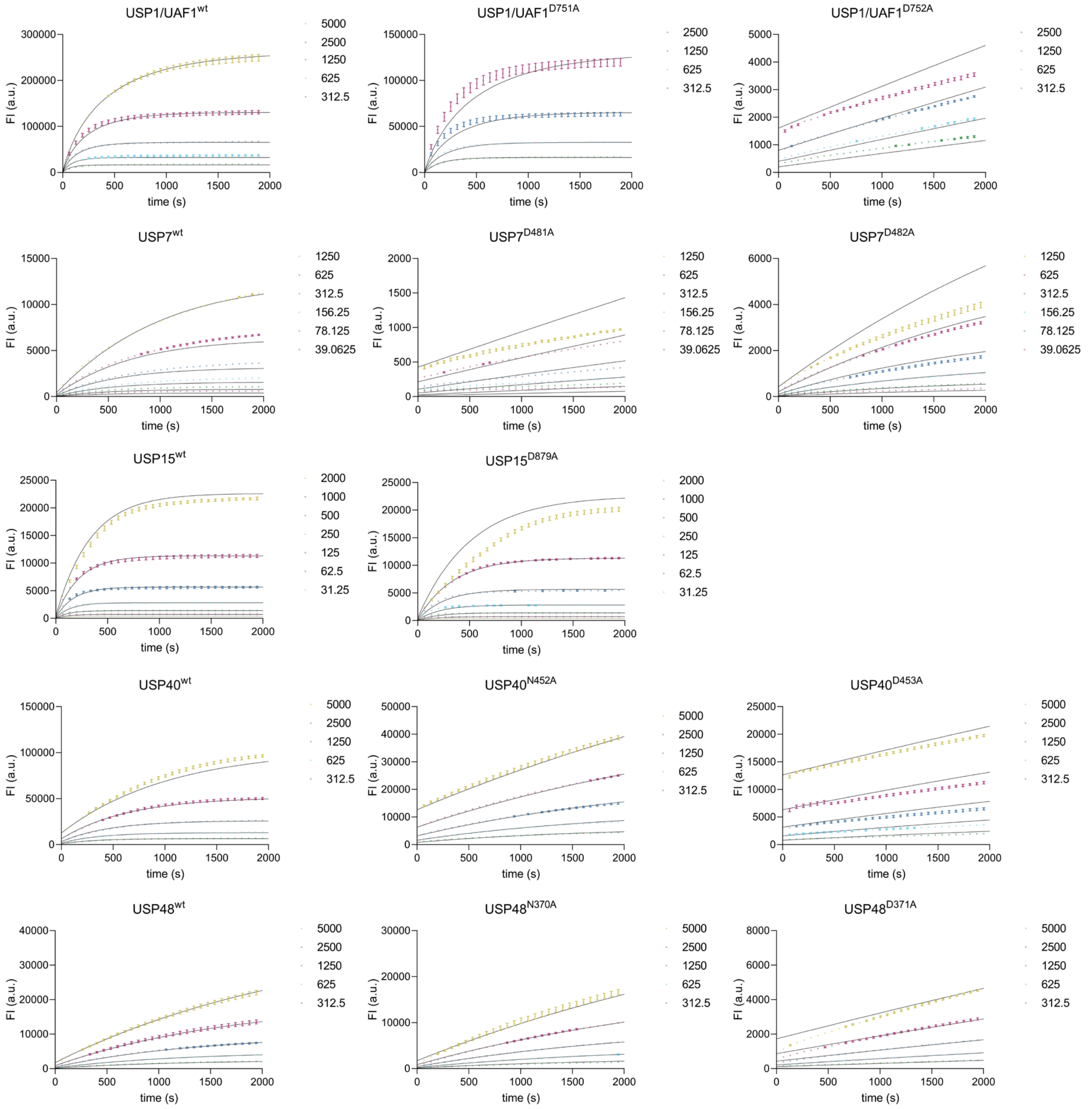

***Supplementary figure 3: Curves generated in Kintek plotted against raw data.*** *Using the method described in Supplementary table 2, we generated curves based on Fluorescence intensity data in order to calculate kinetics (Supplementary table 2). Our kinetic model fits the data well. We got relatively poorer fits to USPs with low activity (USP1*$^{D752A}$*). Original Fluorescence intensity data with standard errors is coloured, Kintek fits are shown in black.*

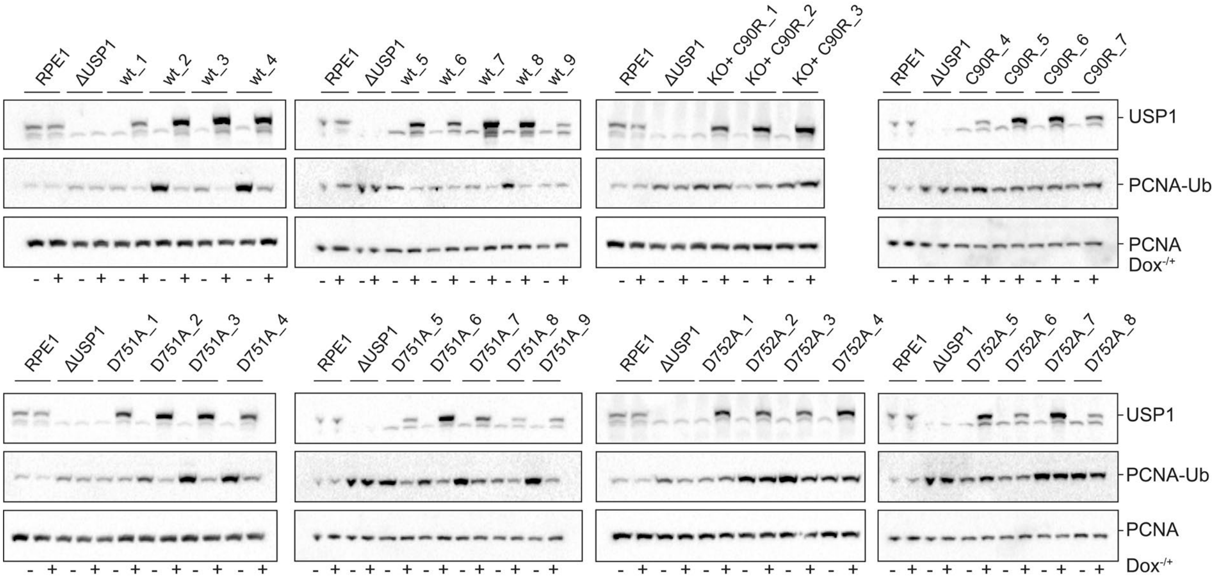

***Supplementary figure 4: Western blot analysis of USP1$^{wt}$, USP1$^{D751A}$, USP$^{D752A}$ USP1$^{C90R}$ used for quantification.*** *RPE1 USP1 knockout cells were complemented with lentiviral expression USP1$^{wt}$, USP1$^{D751A}$, USP1$^{D752A}$ USP1$^{C90R}$. As a control, RPE1 and ΔUSP1 were complemented with an empty vector (EditR). Levels of USP1 expression varied per single clones. Cell lysates before and after doxycyclin induction were stained using antibodies for USP1, PCNA-Ub and PCNA.*

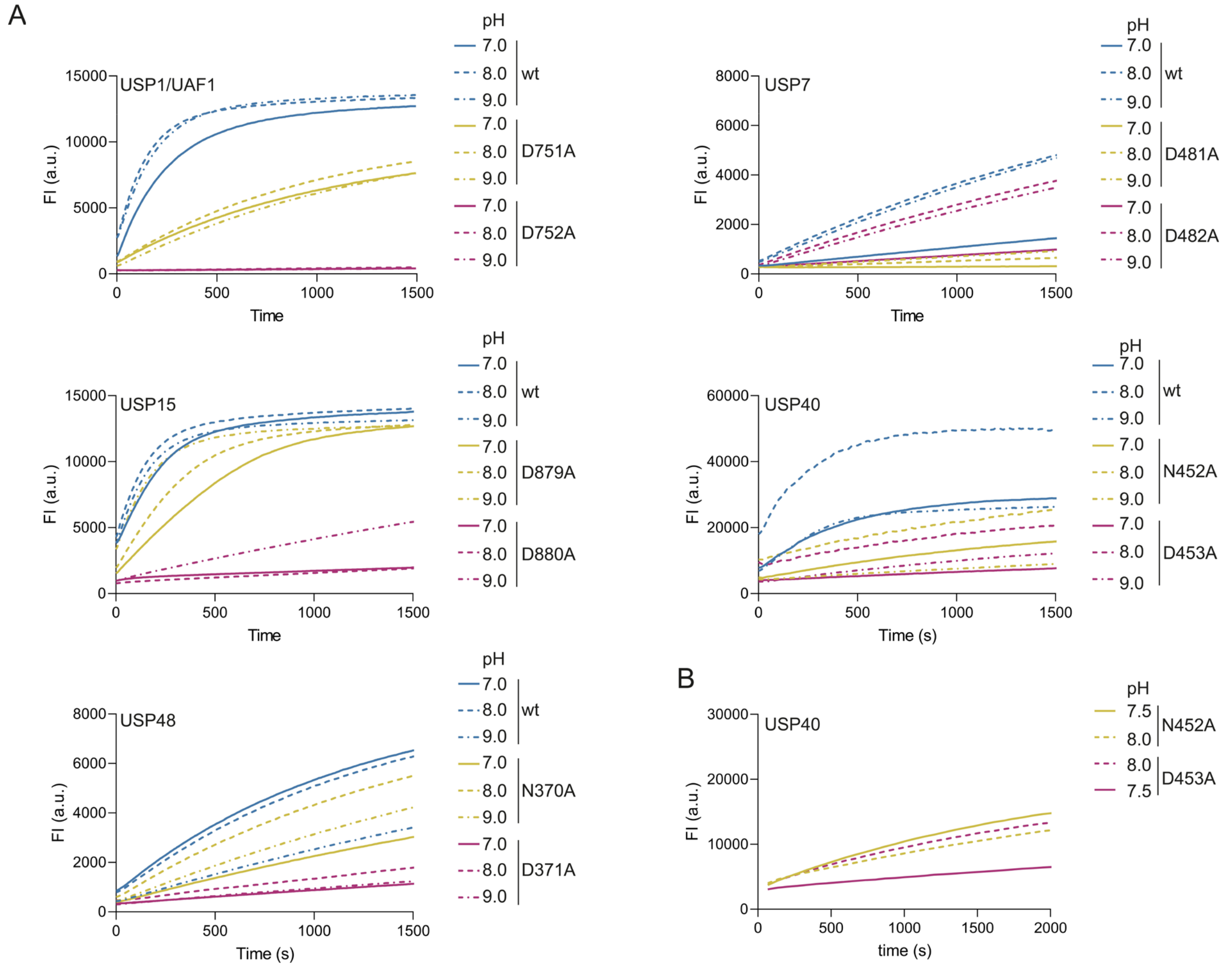

***Supplementary figure 5: USPs do not switch their critical residue at a different pH (A):*** *pH analysis of USPs and their mutants. Single concentrations of enzyme were tested against a single concentration of substrate.* ***(B):*** *Comparison of USP40$^{N452A}$ and USP40$^{D453A}$ at pH 7.5 and pH 8.0. A slight increase in pH boosts USP40$^{D453A}$ activity more than it boosts the activity of USP40$^{N452A}$.*
